# “Generation of two Betacellulin CRISPR-Cas9 knockout hiPSC lines to study affected EGF system paradigm in Schizophrenia”

**DOI:** 10.1101/2025.04.07.647671

**Authors:** Agustin Cota-Coronado, Murray Manning, Dong-Hyung Kim, Joohyung Lee, Andrew Gibbons, Joseph Rosenbluh, Rachel A. Hill, Suresh Sundram

**Affiliations:** Department of Psychiatry, School of Clinical Sciences, Monash University, Clayton, Victoria, 3168, Australia; Cancer Research Program and Department of Biochemistry and Molecular Biology, Biomedicine Discovery Institute, Monash University, Clayton, VIC, Australia; Functional Genomics Platform, Monash University, Clayton, VIC, Australia; Centre for Endocrinology and Metabolism, Hudson Institute of Medical Research, Clayton, Victoria, 3168, Australia; Department of Anatomy and Developmental Biology, Monash University, Clayton, Victoria, 3168, Australia; Mental Health Program, Monash Health, Clayton, Victoria, 3168, Australia

## Abstract

Several members of the epidermal growth factor (EGF) family have been implicated in the biology of schizophrenia (Ketharanathan et al., 2024). The EGF-related ligand, Betacellulin (BTC), plays an important role in the proliferation and differentiation of neural stem cells and our group found markedly reduced BTC levels in patients with schizophrenia. Nevertheless, the interplay of affected BTC and its participation in neural specification and neurodevelopment remains elusive. We generated Knockout (KO) - BTC clones from an existing hiPSC line through CRISPR/Cas9-mediated modification. Furthermore, we validated BTC-KO through genotyping/sequencing, FACS and Western Blot. Finally, we demonstrated trilineage differentiation potential *in vitro*.

## 1. Resource Table

|  |  |
| --- | --- |
| Unique stem cell line identifier | <u>MONPSYi001-A</u><br><u>MONPSYi001-A-1</u><br><u>MONPSYi001-A-2</u> |
| Alternative name(s) of stem cell line | MONPSYi001-A-1<br>MONPSYi001-A-1-BTC-KO<br>MONPSYi001-A-2-BTC-KO |
| Institution | *Monash University* |
| Contact information of the reported cell line distributor | *Agustin Cota-Coronado* |
| Type of cell line | *iPSC* |
| Origin | *Human* |
| Additional origin info<br>(applicable for human ESC or iPSC) | *Age: 55-59*<br>*Sex: Male XY*<br>*Ethnicity: European genetic ancestry* |
| Cell Source | *Fibroblasts, skin tissue* |
| Method of reprogramming | *Sendai virus (non-integrating virus)* |
| Clonality | *Clonal* |
| Evidence of the reprogramming transgene loss (including genomic copy if applicable) | *Described in (Pantazis et al., 2022).* |
| The cell culture system used | *StemFlex serum-free for parental line*<br>*New lines: mTSEr plus and Laminin 521 5ug/mL* |
| Type of the Genetic Modification | *Original line: Gene/mutation Correction CRISPR-Cas9 editing of KOLF2-C1 human iPSCs was performed to correct a 19bp pathogenic deletion in one copy of the ARID2 gene and generated the KOLF2.1J line (Hildebrandt et al, 2019; Pantazis et al, 2022)*<br>*Current line: spCas9-GFP integration with 2 different sgRNAs targeting BTC exon 3* |
| Associated disease | *N/A* |
| Gene/locus modified in the reported transgenic line | *Betacellulin - BTC* |
| Method of modification / user-customisable nucleases (UCN) used, the resource used for design optimisation | *CRISPR/Cas9* |
| User-customisable nuclease (UCN) delivery method | *spCas9 and Lipofectamine* |
| All double-stranded DNA genetic material molecules introduced into the cells | *Recombinant S. pyogenes Cas9-GFP nuclease, purified from an E. coli strain expressing the nuclease. Contains a nuclear localisation sequence (NLS) and C-terminal 6-His tag.*<br>2x synthetic sgRNAs |
| Evidence of the absence of random integration of any plasmids or DS DNA introduced into the cells. | *N/A* |
| Analysis of the nuclease-targeted allele status | *N/A* |
| Homozygous allele status validation | Figure 1B |
| Method of the off-target nuclease activity prediction and surveillance | Cas-OFFinder |
| Descriptive name of the transgene | *N/A* |
| Eukaryotic selective agent resistance cassettes (including inducible, gene/cell type-specific) | *N/A* |
| Inducible/constitutive expression system details | *N/A* |
| Date archived/stock creation date | 05/2024 |
| Cell line repository/bank | MONPSY · Search · hPSCreg |
| Ethical/GMO work approvals | *Project Title: Use and modification of human pluripotent stem cells (hPSCs) to understand the molecular basis of schizophrenia*<br>*Project ID: 38421, 38624*<br>*Chief Investigator: Dr Agustin Cota-Coronado*<br>*Expiry Date: 09/05/2028*<br>*Monash University - MUHREC* |
| Addgene/public access repository<br>recombinant DNA sources' disclaimers<br>(if applicable) | N/A |

## 2. Resource utility

Epidermal growth factor (EGF) is implicated in the pathogenesis of schizophrenia. We identified low Betacellulin concentrations (EGF-related ligand) in people with schizophrenia. We generated novel cell lines to understand the role of BTC in neurodevelopment and the implications of low or absent BTC in Schizophrenia and related psychiatric disorders.

## 3. Resource Details

Schizophrenia is a psychiatric syndrome that is currently diagnosed solely by its clinical features of positive symptoms such as delusions and hallucinations, negative symptoms such as decreased motivation and anhedonia (reduction of pleasure), and cognitive symptoms such as decreased attention and impaired executive function. It has no laboratory or imaging diagnostics, no defining pathology and no known causes or cure. Symptoms and impacts vary widely between individuals with the disorder. Most current treatments are only effective in targeting the psychotic symptoms of schizophrenia with a substantial proportion of people being refractory to treatment. Given that antenatal, early life and adolescent staged environmental factors and high heritability are associated with this adult-onset disorder, neurodevelopmental processes are strongly implicated. Epidermal growth factor (EGF) has been proposed as an important effector molecule in schizophrenia (Ketharanathan et al., 2024; Mostaid et al., 2017; Sotoyama et al., 2013). Additionally, betacellulin (BTC), another ligand member of the EGF family that is capable of signalling through EGF’s receptor, induces neural stem cell (NSC) proliferation in vivo and prevents spontaneous differentiation in culture. Furthermore, it has been shown that *in vivo* infusion in the mouse induced the expansion of neural stem cells and neuroblasts, and promoted neurogenesis in the olfactory bulb and the dentate gyrus (Gómez-Gaviro et al., 2012). Our group recently showed reduced levels of BTC in post-mortem brain tissue from patients with Schizophrenia compared with healthy controls (manuscript in preparation). However, as post-mortem human brain studies do not allow causative determination, we generated these novel resources to understand the role of BTC in early specification and neurodevelopment. We selected the well-characterized hiPSC line KOLF2.1J (Pantazis et al., 2022) and introduced a Cas9-GFP protein through lipofectamine and subsequently two sgRNA directed to the exon 3 in the betacellulin gene. The transfected cells were selected through FACS-sorting for GFP-high cells and performed single-cell deposition. A total of 124 clones were genotyped, of those, 11 clones showed a band in the KO band. Out of those 11 clones, 1 clone did not have a WT band; 2 clones were expanded and sequenced. Subsequently, two clones were genotyped MONPSYi001-A-2-BTC-KO (Hom) was negative for WT PCR, indicating a full KO. Whereas cell line MONPSYi001-A-1-BTC-KO showed a partial deletion of 6 amino acids in the sgRNA1A site, making it a heterozygous clone, compared with the WT line MONPSYi001-A.

The colonies exhibited typical pluripotent stem cell morphology (Fig 1C) under brightfield light. Then, spontaneously differentiated the 3 cell lines and they all showed robust expression of markers belonging to the three germ layers including ectoderm: Nestin and PAX6 (>95% positives) at Day 10 of neural differentiation, mesoderm: α-Smooth Muscle Actin and Brachyury (>90% positives) and endoderm: SOX17 (>90% positives). Furthermore, cells showed reduced BTC at the protein level and were Mycoplasma negative, validated with nested PCR (Supplementary Figure 1). To our knowledge, these are the first hiPSC lines that can test the low BTC paradigm schizophrenia representing valuable resources for disease modelling and drug screening based on clinical data.

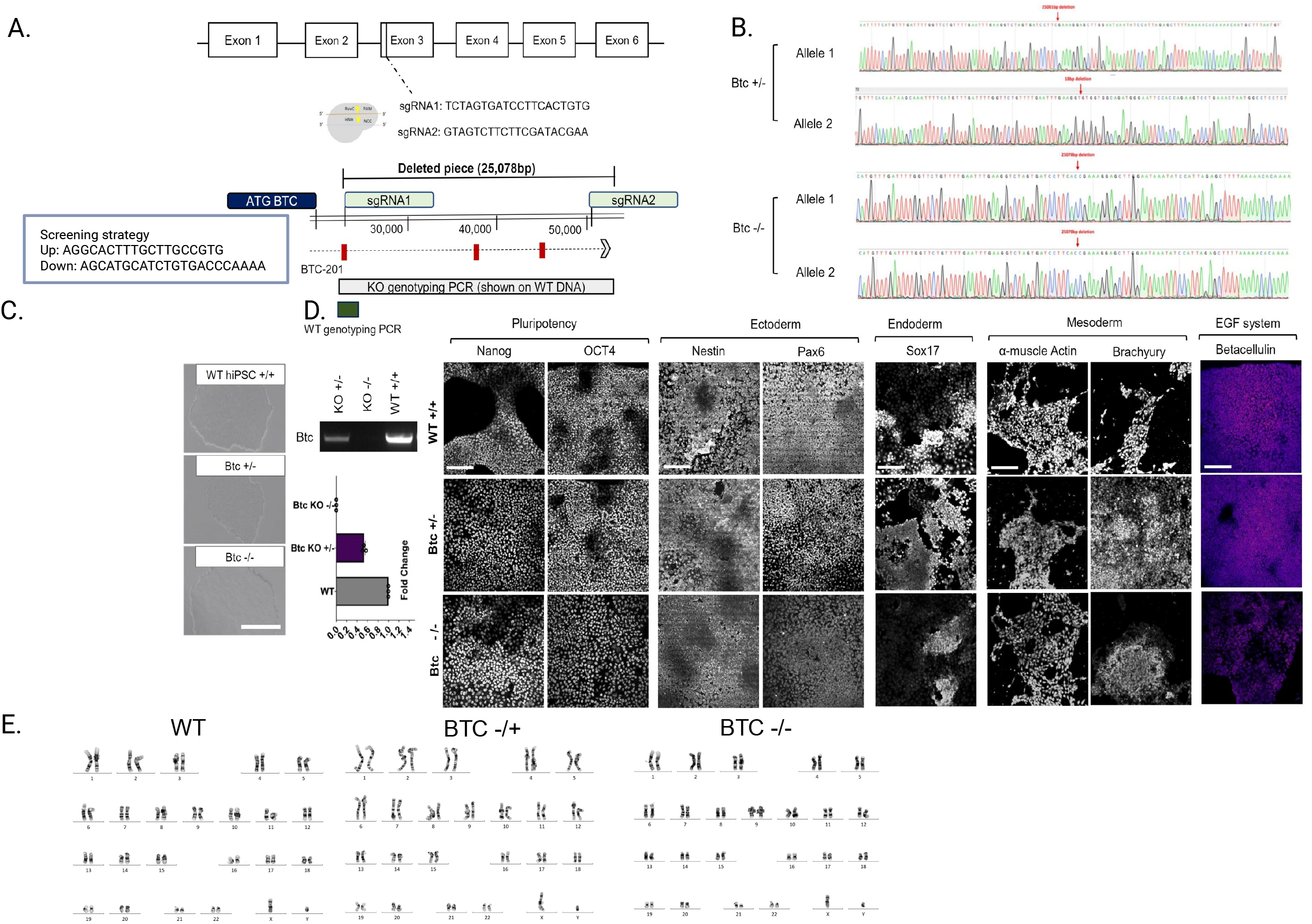

## 4. Materials and Methods

### 4.1 hiPSC culturing and passaging

KOLF2.1J human induced pluripotent stem cell line (The Jackson Laboratory, cat # JIPSC1000) and isogenic clones derived from this line were grown in mTSER Plus feeder-free media (STEMCELL, cat # 85850). 10uM Rock inhibitor (STEMCELL, Cat # 72304) was added to mTSER Plus media for 18–24 h after splitting and thawing.

### 4.2 CRISPR-Cas9 knockouts

KOLF2.1J line was transfected with a high-fidelity Cas9-eGFP using Lipofectamine STEM (Thermo Fisher, cat # STEM00001). A Recombinant S. pyogenes Cas9 nuclease, purified from an E. coli strain expressing the nuclease. It holds a nuclear localisation sequence (NLS) and C-terminal 6-His tag at 10 µg/µL. Combined Solution I: 25 UG Cas9-GFP, 3.2µg sgRNA1A, 3.2µg sgRNA2and 500µl OPTI-MEM was used. Solution II: 40µl Lipofectamine STEM and 500µl OPTI-MEM were incubated at RT for 10min, then added to a 2 x T25 flask with ∼70% confluent cells 2 days after transfections the cells were FACS sorted for GFP high cells into 12x 96-well plates (single cell deposition). Bulk cells were collected for gDNA extraction and PCR.

### 4.3 Genotyping

KO PCR products were sequenced to verify the exact KO mutations for the transfected 2 clones. The cell line MONPSYi001-A-2-BTC-KO (Homozygous) was negative for WT PCR, indicating a full KO. Whereas cell line MONPSYi001-A-1-BTC-KO showed a partial deletion of 6 amino acids in the sgRNA1A site, making it a heterozygous clone.

### 4.3 Western Blot

20 µg of protein was run on either 8 or 12% acrylamide gel and transferred to a polyvinylidene difluoride (PVDF) membrane using the Trans-Blot Turbo RTA Transfer Kit (Bio-Rad, Cat # 1704274). Primary antibodies raised against BTC (1:500, Invitrogen, Cat # PA5-102636), Cas9 (1:500, Invitrogen, Cat # MA1-201), β-actin (1:1000, Sigma-Aldrich, Cat # A5316), were applied overnight at 4°C and labelled with AlexaFluor488 anti-mouse (1:1000, Invitrogen, Cat # A21202) or AlexaFluor647 anti-rabbit (1:1000, Invitrogen, Cat # A31573) secondary antibodies for 1h at room temperature. PVDF membranes were scanned and imaged using the ChemicDoc MP Imaging System (Bio-Rad). BTC was normalized against the internal control, β-actin, and Cas9 were normalized against the internal control, β-tubulin. Blot density was analyzed using the Fiji-ImageJ v1.54f software with the gel analysis routine method.

### 4.4 Mycoplasma detection

Two independent nested PCR-based protocols (I/II) were performed for detecting Mycoplasma/Acholeplasma species. Two positive controls: +ve1 and +ve2 are shown as Mycoplasma positive samples, giving the expected products at 219-426bp (protocol I) and 165-355bp (protocol II). None of the new cell lines gave a positive band, indicating them as mycoplasma-free.

### 4.5 Pluripotency assessment

iPSC clones were grown on laminin-521 coated Ibidi 8-well plates (cat# fixed with 4% PFA, and stained with pluripotency markers listed in Table 3. Images were taken on a Zeiss 880 confocal microscope. The percentage of cells positive for OCT3/4 and NANOG was calculated from immunofluorescence images using the Fiji software, compared with total DAPI-positive cells.

### 4.6 Differentiation potential

iPSC clones were differentiated into three layers according to STEMdiff™ Trilineage Differentiation Kit (StemCell, Cat# 05230), fixed in 4% PFA, and stained with differentiation markers listed in Table 3. Images were taken on a Nikon C1 confocal microscope at 20X magnification (Fig 1F).

---

**Table 1:** Characterization and validation.

| **Classification**<br>(optional *italicized*) | **Output type** | **Result** | **Data** |
| --- | --- | --- | --- |
| **Schematic of a transgene/genetic modification** | Schematic illustrating the structure and location of the introduced genetic modification | *A visual representation of edited allele, showing exon structure or protein product; for transgene showing important functional elements.* | *Position of editing event, gRNA in the gene/on the chromosome, Sanger sequencing track(s) illustrating allelic status), suggested for Fig 1 panel A* |
| **Morphology** | Photography<br>*[mandatory]* | *All iPSC lines are morphological normal* | *Bright-Field image Fig 1A* |
| **Pluripotency status evidence for the described cell line** | Qualitative analysis (*i.e. Immunocytochemistry, western blotting*)<br>*[mandatory]* | *Assess staining/expression of pluripotency markers: OCT3/4*<br>*NANOG* | *Figure 1D* |
|  | Quantitative analysis (*i.e. Flow cytometry, RT-qPCR*) of expression alongside well-known/published hPSC line *[mandatory]* | *Assess % of positive cells by confocal microscopy (cell counting)*<br>*OCT3/4: Red > 95%*<br>*NANOG: Purple > 95%* | *In request to the authors* |
| **Karyotype** | Karyotype (G-banding *[mandatory]*) and higher-resolution, array-based assays (KaryoStat, SNP, etc.) | *46XY,*<br>*Resolution 450 bhps*<br>*G-banding* | *Figure 1D* |
| **Genotyping for the desired genomic alteration/allelic status of the gene of interest** | PCR across the edited site or targeted allele-specific PCR<br>*[mandatory]* | *Sequencing of edited part in exon 3* | *Figure 1B* |
|  | Evaluation of the - (homo-/hetero-/hemi-) zygous status of introduced genomic alteration(s) | *Sequencing of edited part in exon 3* | *Figure 1B* |
|  | Transgene-specific PCR (when applicable) | *N/A* | *N/A* |
| **Verification of the absence of random plasmid integration events** | PCR/Southern<br>*[mandatory]* | *Not performed* | *N/A* |
| **Parental and modified cell line genetic identity evidence** | STR analysis, microsatellite PCR (mPCR) or specific (mutant) allele seq | *DNA Profiling* | *Figure 1B* |
|  | *[mandatory]* | *e.g. Include minimum amelogenin+ 8 loci (D5S818, D13S317, D7S820, D16S539, vWA, Th01, TPOX, CSF1PO) and specify if matched or not. Tabulate the STR results in an excel file* | *N/A* |
| **Mutagenesis / genetic modification outcome analysis** | Sequencing (genomic DNA PCR or RT-PCR product)<br>*[mandatory]* | *RT-PCR product of BTC-KO Het and Homo* | *Figure 1C* |
|  | PCR-based analyses | *N/A* | *N/A* |
|  | Southern Blot or WGS; western blotting (for knock-outs, KOs) | *In request to the authors* | *N/A* |
| **Off-target nuclease activity analysis** | PCR across top 5/10 predicted top likely off-target sites, whole genome/exome sequencing<br>*[Optional but highly recommended if Cas editing is used]* | *Not performed* | *N/A* |
| **Specific pathogen-free status** | Mycoplasma<br>*[Mandatory]* | *Mycoplasma negative in all clones. Nested PCR* | *Supplementary Fig 1* |
| **Multilineage differentiation potential** | e.g. Embryoid body formation OR Teratoma formation OR Scorecard OR Directed differentiation<br>*[Mandatory]* | *The STEMdiff™ Trilineage Differentiation Kit (StemCell) was used to test differentiation potential. We confirmed the expression of endoderm (SOX-17), mesoderm (<-SMA, Brachyury) and ectoderm (NESTIN and PAX6) markers in all clones.* | *IF proof of marker expression in each germ layer* |
| **List of recommended germ layer markers** | Expression of these markers has to be demonstrated at mRNA (RT PCR) or protein (IF) levels, at least 2 markers need to be shown per germ layer | *Ectoderm: PAX6/NESTIN*<br>*TUJ1, MAP2, DCX and Ki67*<br>*Endoderm: SOX17*<br>*Mesoderm: BRACHYURY and alpha-Smooth Muscle Actin* | *IF with specific antibodies* |
| **Outcomes of gene editing experiment (OPTIONAL)** | Brief description of the outcomes in terms of clones generated/establishment approach/screening outcomes | *Individual clones screened: 102*<br>*Clones + for KO PCR: 11*<br>*Clones negative for WT PCR : 1*<br>*Clones expanded: 3* | *In request to the authors* |
| **Donor screening (OPTIONAL)** | HIV 1 + 2 Hepatitis B, Hepatitis C | *Not performed* | N/A |
| **Genotype - additional histocompatibility info (OPTIONAL)** | Blood group genotyping | *Not performed* | N/A |
|  | HLA tissue typing | *Not performed* | N/A |

**Table 2:**
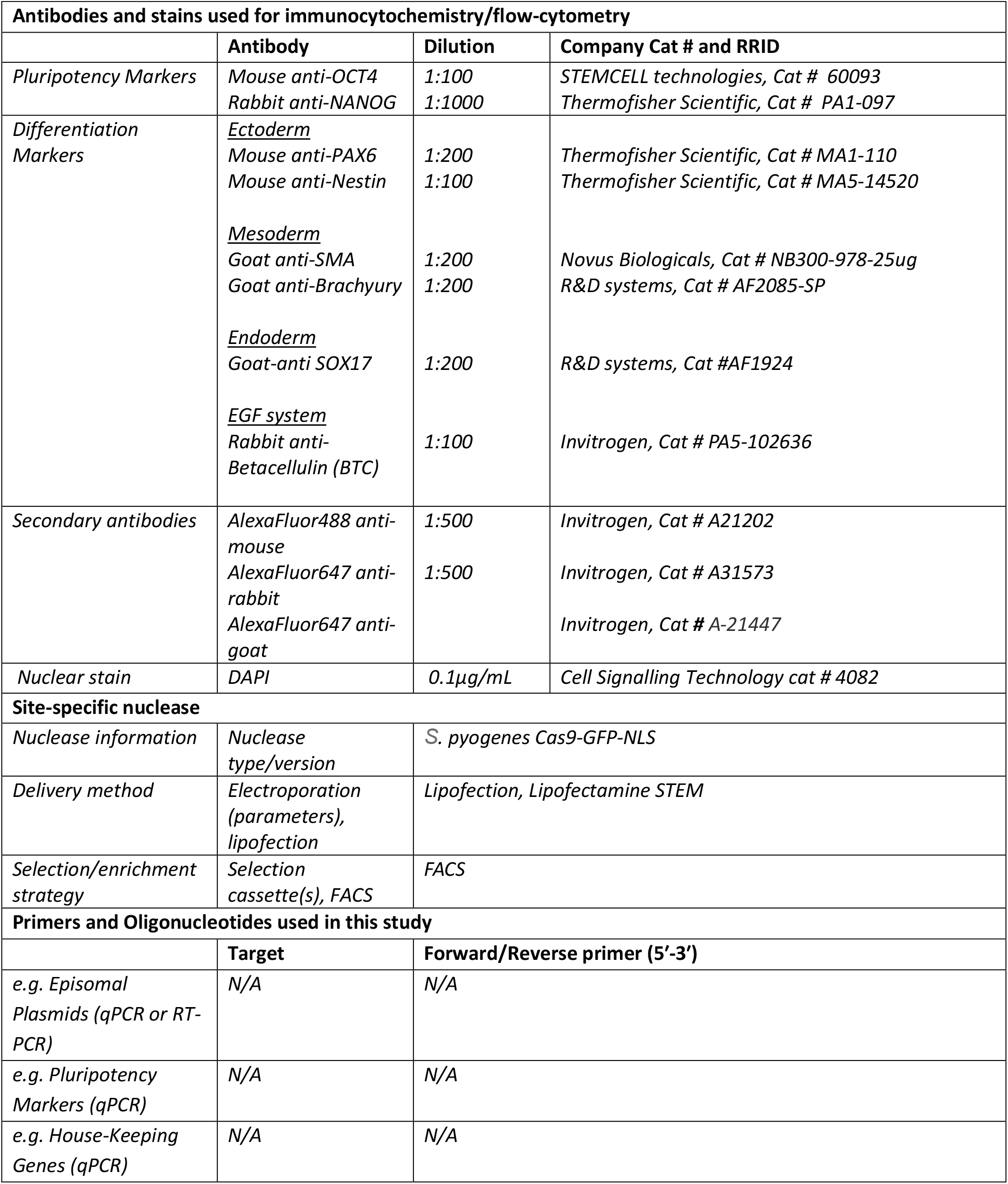
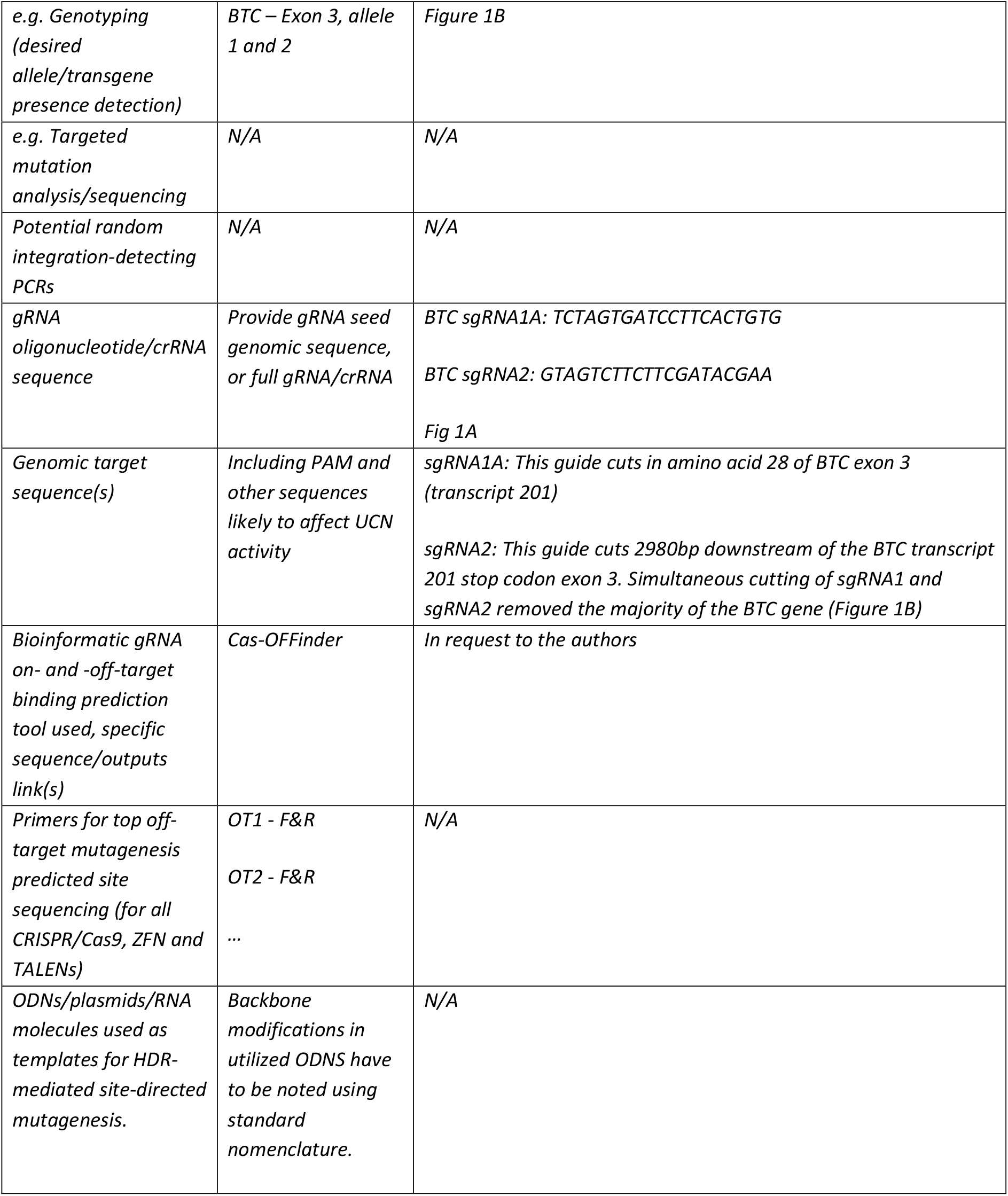
Reagents details.

