## Supplementary figures and images for "“Generation of two Betacellulin CRISPR-Cas9 knockout hiPSC lines to study affected EGF system paradigm in Schizophrenia”"

### Supplemental Figure 1

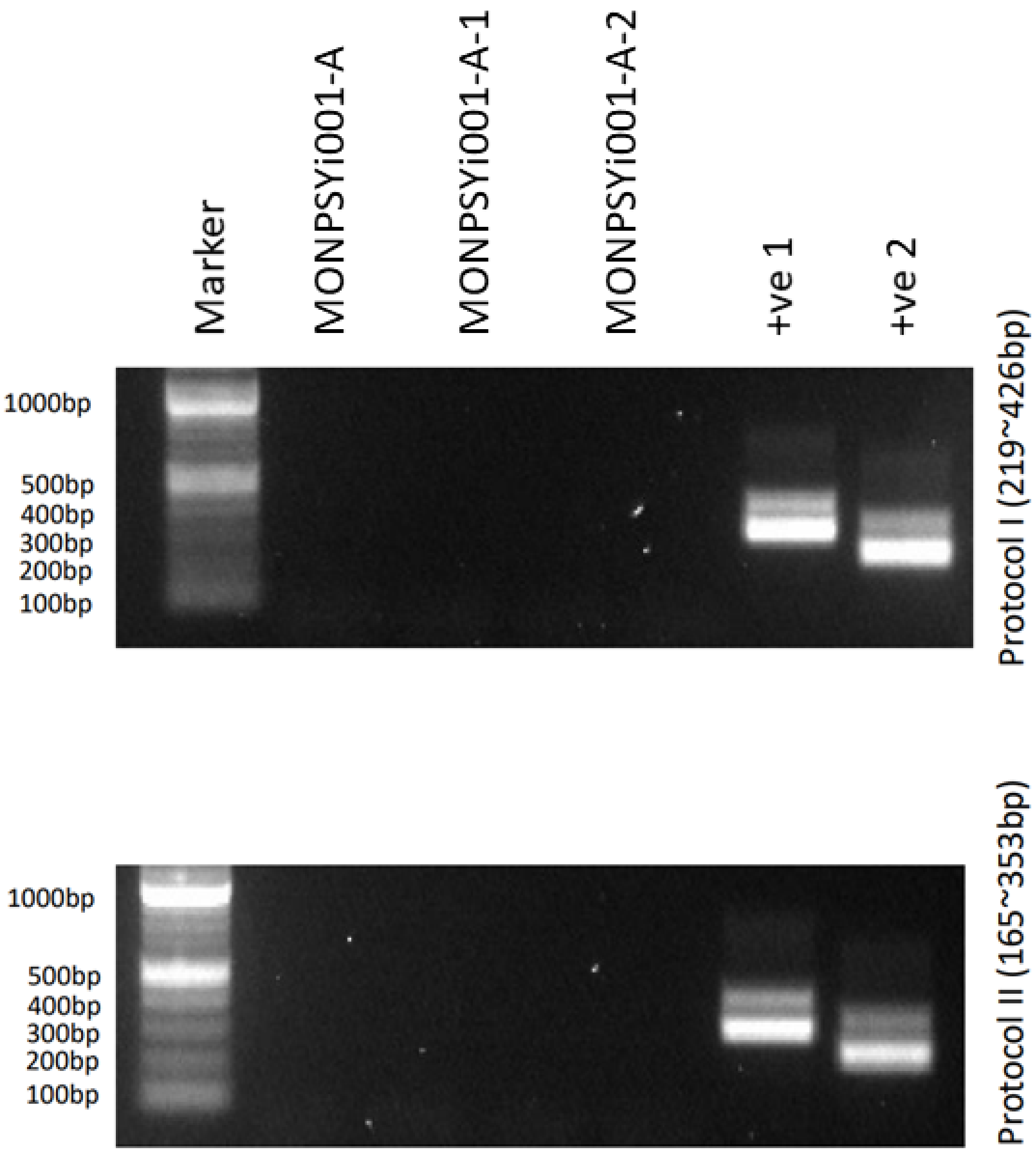
